# Legacy of the lost and pressure of the present: Malagasy plant seeds retain megafauna dispersal signatures but downsize under human footprint

**DOI:** 10.1101/2025.04.16.649107

**Authors:** Yuanshu Pu, Alexander Zizka, Renske E. Onstein

**Affiliations:** German Centre for Integrative Biodiversity Research (iDiv) Halle-Jena-Leipzig, Leipzig, DE; Univeristy of Leipzig, Leipzig, DE; Department of Biology, Philipps-University Marburg, Marburg, DE; Naturalis Biodiversity Centre, Leiden, NL

**Keywords:** defaunation, endozoochory, frugivory, Madagascar, megafauna extinction, seed dispersal, structural equation model, trait matching

## Abstract

Past megafaunal extinctions and ongoing declines of remaining large-bodied frugivores limit the dispersal of large-seeded plants, contributing to their (local) demise, thereby ‘downsizing’ plant seeds across assemblages. Here, we test this hypothesis by integrating trait and distribution data for 2,852 endozoochorous plant species, 48 extant and 15 extinct frugivore species across 361 assemblages on Madagascar. Using structural equation models, we show that mega-seeded plants – with seeds that are too large to be swallowed by any extant Malagasy frugivore – have larger seeds in assemblages where past megafrugivores were larger, highlighting the legacy of past interactions on persisting mega-seeded plants. Furthermore, human footprint reduces seed sizes directly and indirectly by downsizing of extant frugivore in the assemblages. Such human-driven seed downsizing can lead to erosions of important ecosystem functions such as tropical forest carbon storage.

## Introduction

Co-evolution and niche tracking between mutualistic plants and vertebrates shape the composition of contemporary local assemblages and their interaction networks (Chagnon et al., 2015; Dalsgaard et al., 2021; Durand-Bessart et al., 2023). Additionally, the impact of climate and human activities on the spatial distribution of species modifies assemblage compositions (Barnagaud et al., 2017; Joswig et al., 2021; Martins et al., 2021). However, the respective magnitude of past and present mutualistic interactions and how they have shaped contemporary assemblage trait structures remain unclear.

Endozoochory is the mutualism between fruit-producing plants and fruit-eating animals (frugivores, hereafter), in which the frugivores consume fleshy fruits and disperse the contained seeds after gut passage. In this mutualism, the size of the ingested seeds is constrained by the frugivore’s gape width, which aligns with the frugivore’s body mass (Burns, 2013; Lord, 2004; Wheelwright, 1985). Larger seeds benefit from dispersal by larger-bodied animals, with dispersal over longer distances, higher seedling survival rates, reduced resource competition with relatives, and increased opportunities for successful colonization of new habitats (Chapman, 2005; Onstein et al., 2019; Pires et al., 2018; Wotton & Kelly, 2012). However, larger-seeded plants are restricted to fewer frugivore species that can ingest their seeds. Indeed, spatial patterns of ‘trait matching’ (i.e., a positive correlation) between frugivore (e.g., gape width or body mass) and plant (e.g., fruit width or seed width) have been discovered worldwide (Dehling et al., 2014; Mack, 1993; McFadden et al., 2022). However, inconsistencies in such spatial correlation can reveal mutualism ‘disruption’ (Lim et al., 2020; Méndez et al., 2022), as extinctions of large-bodied animals reduce the maximum body size in many assemblages worldwide (i.e., ‘downsizing’; Dirzo et al., 2014; Young et al., 2016). Consequentially, seed dispersal limitation following the frugivore downsizing can lead to structural (e.g., secondary plant extinctions) and functional (e.g., loss of long distance dispersal) decay in plant assemblages (Donoso et al., 2020; Fricke et al., 2022). However, it remains unclear whether it could ultimately lead to changes in assemblage-wide trait structures, such as seed size.

Frugivore downsizing has been particularly severe in Madagascar, where all animals with a body mass over 10 kg rapidly went extinct ca. 1,000 years ago (Crowley, 2010). Additionally, 94% of the remaining lemur species, the largest and most important extant frugivore guild in Madagascar, are threatened with extinction (Albert-Daviaud et al., 2018; IUCN, 2022; Razafindratsima, 2015). As more than 30% of plant species in Madagascar depend on endozoochorous seed dispersal (3,324 out of 9,434 endemic species; Albert-Daviaud et al., 2018; Antonelli et al., 2022), the loss of frugivorous lemurs may have detrimental consequences for the Malagasy flora. Interestingly, the seeds of some plant species, e.g., *Borassus madagascariensis* (Arecaceae, seed width 67.50 mm) and *Tsebona macrantha* (Sapotaceae, seed width 37.50 mm), are larger than the gape width of any extant frugivorous lemur. The largest seeds recorded to be dispersed by extant frugivores are 24.6 mm wide, from *Canarium madagascariense* (Burseraceae; Albert-Daviaud et al., 2020; Razafindratsima & Martinez, 2012), suggesting that plants with seeds larger than this threshold are not effectively dispersed through endozoochory anymore. These species (mega-seeded plants, hereafter) are therefore ‘anachronistic’ – i.e., they possess large fruits and seeds that seem adapted to past biotic interactions but maladapted in current ecosystems (Janzen & Martin, 1982). It is unclear how these mega-seeded plants have persisted in ecosystems since the extinction of their primary megafaunal seed dispersers (megafrugivores, hereafter), and whether they hold signatures of past trait matching in their distribution range (Heinen et al., 2023; Méndez et al., 2022).

In addition to seed dispersal limitation due to human-driven frugivore downsizing, endozoochorous plants, particularly those with large seeds, also face direct human impact. Due to allometric scaling, large seeds are often found in large trees (Leishman et al., 1995; Moles et al., 2005), which are especially susceptible to human activities such as timber logging (Asner et al., 2009). In Madagascar, over 44% of the forest has been cleared since the 1950s by humans, for instance for slash-and-burn agriculture and pasture (Scales, 2011; Vieilledent et al., 2018). Moreover, the slow growth and recruitment rates of large trees with large seeds further constrain their recovery post disturbance (Barczyk et al., 2024; Osuri & Sankaran, 2016; Palma et al., 2021). Despite these theoretical expectations, it remains unknown whether human footprint has affected the distribution of larger-seeded plants in Madagascar.

In this study, we take an assemblage-wide approach to disentangle the direct and indirect effects of human footprint on seed size in endozoochorous plants in Madagascar, considering the contemporary and past distributions of (mega-)frugivores. First, we hypothesize (H1) that human footprint leads to downsizing of seeds in assemblages, both directly (e.g., due to selective logging of large-seeded trees) and indirectly, via downsizing of extant frugivores (e.g., due to hunting; ‘downsizing-effect hypothesis’). Accordingly, we expect that maximum frugivore and seed sizes in assemblages decrease with increasing human footprint, and that there is a positive relationship between frugivore body mass and seed size across assemblages (reflecting spatial trait matching). Second, we hypothesize (H2) that the occurrence of anachronistic mega-seeded plant is spatially matched with the previous occurrence of now extinct megafrugivores, but not with the occurrence of extant frugivores (‘legacy-effect hypothesis’). Hence, we expect that among assemblages with mega-seeded plants, seeds are larger in assemblages where larger (now extinct) megafrugivores occurred. In contrast, among assemblages without mega-seeded plants, we expect that seeds are larger in assemblages with larger extant frugivores. By testing these hypotheses, we provide insights into how past megafrugivores and present human impact jointly shape contemporary seed size distributions, with broader implications for the potential erosion of ecosystem functions due to seed downsizing in tropical forests.

## Material and methods

### Endozoochorous plant species

We used the species checklist of endemic endozoochorous seed plants in Madagascar from Albert-Daviaud et al. (2020; Table S1). Endozoochory was defined by the presence of edible rewards such as fruit pulp or an aril (van der Pijl, 1972). Out of 3,241 endozoochorous plant species, we obtained occurrence data for 2,852 species from Ralimanana et al. (2022), based on both validated records from the iNaturalist research-grade observations and specimens from the Missouri Botanical Garden, accessed via the Global Biodiversity Information Facility (GBIF.org, 2020a, b). The raw records were filtered by excluding records from centroids of Madagascar and its provinces, the capital city Antananarivo, and registered research institutions, using *CoordinateCleaner* (Zizka et al., 2019).

We obtained seed widths (the second largest dimension of a seed, which is the limiting axis for animals to ingest a seed) for all endemic endozoochorous plants in Madagascar from Albert-Daviaud et al. (2020) (Table S1). Seed widths of 1,323 out of 3,241 species were collected from botanical descriptions, databases, specimens from the Royal Botanic Gardens, Kew and the Muséum National d’Histoire Naturelle, Paris (Albert-Daviaud et al., 2020). Seed widths of the remaining species were imputed using multivariate imputation by chained equations (MICE; van Buuren & Groothuis-Oudshoorn, 2011), based on eight traits related to fruit or seed morphology, life form, bioclimate and vegetation formation of 8,788 species across the Malagasy flora (Albert-Daviaud et al., 2020). Finally, among all endemic endozoochorous plants in Madagascar, 15 species had seeds that were wider than 24.6 mm, which was the width of the largest seed (from *Canarium madagascariense*) recorded to be dispersed by any extant Malagasy frugivore (Albert-Daviaud et al., 2020; Razafindratsima & Martinez, 2012). These species were considered ‘anachronistic’ and classified as mega-seeded plants in our analyses.

### Extant and extinct frugivores

We considered all extant vertebrate species on Madagascar whose diets consist for at least 50% of fruits as ‘frugivores’ (48 species in total). We used this threshold because the distribution and trait (i.e., body mass) of these animals were expected to be closely aligned with the distribution and trait (i.e., seed width) of their endozoochorous food plants, due to strict mutualistic dependence (Eppley et al., 2020). For extant frugivores, we used the frugivory index data from EltonTraits 1.0 (Wilman et al., 2014) and Galán-Acedo et al. (2019). 16 species had no dietary data available, for which we inferred the most probable dietary group based on information from other species within the same genus. The final dataset included 28 primates, 3 bats, 11 birds and 6 rodents (Table S2). We obtained distribution ranges for these species from the International Union for Conservation of Nature Red List, ver. 2022-2 (IUCN, 2022), and body mass data by calculating the average values across available species-level data from Wilman et al. (2014), Galán-Acedo et al. (2019) and Razafindratsima et al. (2018; Table S2).

As extinct megafrugivores in Madagascar, we included four elephant bird species, nine giant lemur species and two giant tortoise species, following Méndez, et al. (2022, 2024; Table S2). We obtained historical distribution maps for these species from Méndez et al. (2022) and Pedrono et al. (2013), which were reconstructed based on fossil/subfossil sites and distributions of coeval species (i.e., species that existed at the same time), using MInOSSE (Carotenuto et al., 2020). We obtained body mass data of extinct megafrugivores from Jungers et al. (2008), Hansford and Turvey (2018), Hansen et al. (2008) and the Turtle Taxonomy Working Group (2021), where body masses were inferred from bone and carapace measurements (Table S2).

### Assemblage-level data

We rasterized the terrestrial area of Madagascar into 649 grid cells of 30x30 km, each representing an assemblage of co-occurring endozoochorous plant and frugivore species (Table S3). By using this approach, we aimed to balance between spatial resolution and data availability. First, occurrence and range maps were intersected with the 649 assemblages to generate a presence-absence matrix for all plant and frugivore species. Then, we calculated the 95^th^ percentile maximum plant seed width (maximum seed width, hereafter) and the 95^th^ percentile maximum frugivore body mass (maximum body mass, hereafter) for each assemblage. By using this measure instead of maximum values, we reduced the influence of extreme outliers and thereby better captured the upper range of plant seed widths and frugivore body masses in an assemblage.

We excluded assemblages on the coast with less than 50% terrestrial area, assemblages that lacked extinct frugivores, assemblages with fewer than three extant frugivore species, and assemblages with fewer than three endozoochorous plant species (361 out of 649 assemblages remained). This allowed us to focus on patterns that emerged from mutualistic interaction-related processes as outlined in our hypotheses, while avoiding biased results due to outlier assemblages with few species.

### Human footprint and environmental factors

We obtained a human footprint map of 2009 from Venter et al. (2016), which compiled and quantified different human stressors related to land use transformation (built environment, crop and pasture lands, night-time lights), transportation infrastructure (roads, railways and navigable waterways) and human population density. We calculated the average value of the cumulative human footprint for each assemblage in Madagascar.

While our focus was on the effects of human footprint and frugivore body mass on seed width, we accounted for environmental factors to avoid confounding effects. Environmental conditions, such as higher temperature and precipitation, can independently promote larger seeds by providing greater energy and resource availability (Moles et al., 2007), or promote smaller mammalian frugivores due to heat conservation mechanisms (Rodríguez et al., 2008). Without considering these factors, the relationship between seed width and frugivore body mass could be absent due to confounding environmental effects, or conversely, observed relationships could be misattributed to mutualistic interactions rather than shared environmental drivers. We obtained environmental data from MadaClim (https://madaclim.cirad.fr/). For each assemblage, we calculated mean values of 26 variables related to temperature, precipitation, topography, solar radiation, water deficiency and percentage of forest cover (for a complete list of variables, see Table S4). Then, to reduce the dimensionality of the dataset while capturing the most variance, we normalised and scaled all variables and conducted principal component analysis (PCA). The first two principal components (PC) were used in subsequent analyses.

### Structural Equation Modelling

To disentangle the direct and indirect causal effects of frugivores and human footprint on the distribution of plant seed widths in Madagascar, we used structural equation models (SEM) as implemented in the R package “lavaan” v0.6-12 (Rosseel, 2012) in R 4.4.1 (R Core Team, 2024). We defined the *a priori* model structure as shown in Fig. S1. First, the model included the ‘downsizing’ effect of human footprint on maximum plant seed width, directly and indirectly by affecting maximum extant frugivore body mass. Moreover, the ‘legacy’ effect of extinct megafrugivores was modelled by testing for a relationship between assemblage maximum extinct frugivore body mass and maximum plant seed width. Furthermore, we included the effects of environmental PC1 and PC2 on plant seed width, extant and extinct frugivore body mass, and human footprint. We allowed covariance between human footprint and maximum extinct frugivore body mass, and covariance between maximum body masses of extant and extinct frugivores. By allowing the residual variances of two observed variables to be correlated, we acknowledge that these variables may share underlying historical or environmental drivers that are not explicitly included in the model. We log-transformed biological variables (i.e., seed widths and body masses) to approach normal distributions of model residuals, and scaled all variables to 0-1 to compare standardised effect sizes among predictor variables. We used the “MLR” estimator when conducting the SEM, which introduces data-based corrections to the test statistics and standard errors to offset the bias introduced by the non-normal distribution of model residuals (Rosseel, 2012).

We fitted three SEMs to the data. First, a SEM using all 361 assemblages to identify direct and indirect effects of human footprint on seed size (H1). Then, to evaluate the effect of past interactions on seed size of anachronistic mega-seeded plants (H2), we used a comparative framework, in which we compared the effects of the predictor variables for mega-seeded and non-mega-seeded plants. To this end, we fitted the same *a priori* SEM with two subsets of the data: assemblages with mega-seeded plants (66 assemblages), and assemblages without mega-seeded plants (295 assemblages) (Fig. S2).

To refine the model structure, we followed a stepwise removal of statistically non-significant paths starting from the *a priori* model. We excluded the path with the highest p-value from the model, reran the SEM, and repeated this procedure until only paths representing a significant effect remained. Following Hooper et al. (2008), since different indices capture different aspects of model fit, we considered a final model a good fit if a combination of conditions was met: the root mean square error of approximation (RMSEA) smaller than 0.05, the comparative fit index (CFI) greater than 0.95, and the Chi-Square test not significant (p > 0.05).

### Randomisations and spatial autocorrelation

To assure that relationships revealed in the final SEM models deviated from random expectations, we reran the final SEM after randomizing maximum seed widths across assemblages, and repeated this 1,000 times. We obtained p-values and effect sizes for each predictor of maximum seed width in the randomized scenarios, and assessed whether observed effect sizes and p-values from the empirical data significantly differed (i.e., outside 95% distribution) from randomized inferences.

Significant effects recovered from the SEM analyses may be derived from spatial autocorrelation among assemblages. Therefore, we tested for spatial autocorrelation in our focal variable – maximum seed width – and conducted spatial autoregressive (SAR) modelling to correct for potential effects of spatial autocorrelation on outcomes. We used the R package “spdep” v1.2-5 (Bivand & Wong, 2018). First, we defined the spatial weight matrix for the SAR models, using a minimum distance that can link each assemblage grid to at least one other assemblage grid (i.e., 44.6 km). Next, we evaluated the estimate and significance of all direct predictor variables on maximum seed widths that were identified as significant in the SEM, in a non-spatial linear regression model and a SAR model, which were directly comparable, as they only differed in their correction for spatial autocorrelation. Finally, we calculated Moran’s I values for the linear model, the residuals of the linear model, and the SAR model, to evaluate the role of spatial autocorrelation in our data and model outcomes.

## Results

Maximum seed widths, maximum extant frugivore body masses, maximum extinct frugivore body masses, and human footprint, were distinctly distributed across Madagascar. Assemblages in the eastern rainforests and the western drylands had larger maximum seed widths than assemblages in the central highlands and the southern spiny thickets (Fig. 1a). Extant frugivores with the largest body masses were concentrated in the eastern rainforests (Fig. 1b), whereas extinct frugivores with the largest body masses were primarily distributed in the dry west (Fig. 1c). Finally, human footprint was highest in the central highlands around major cities such as Antananarivo and Fianarantsoa, as well as in southern and eastern coastal areas (Fig. 1d).

**Figure 1:**
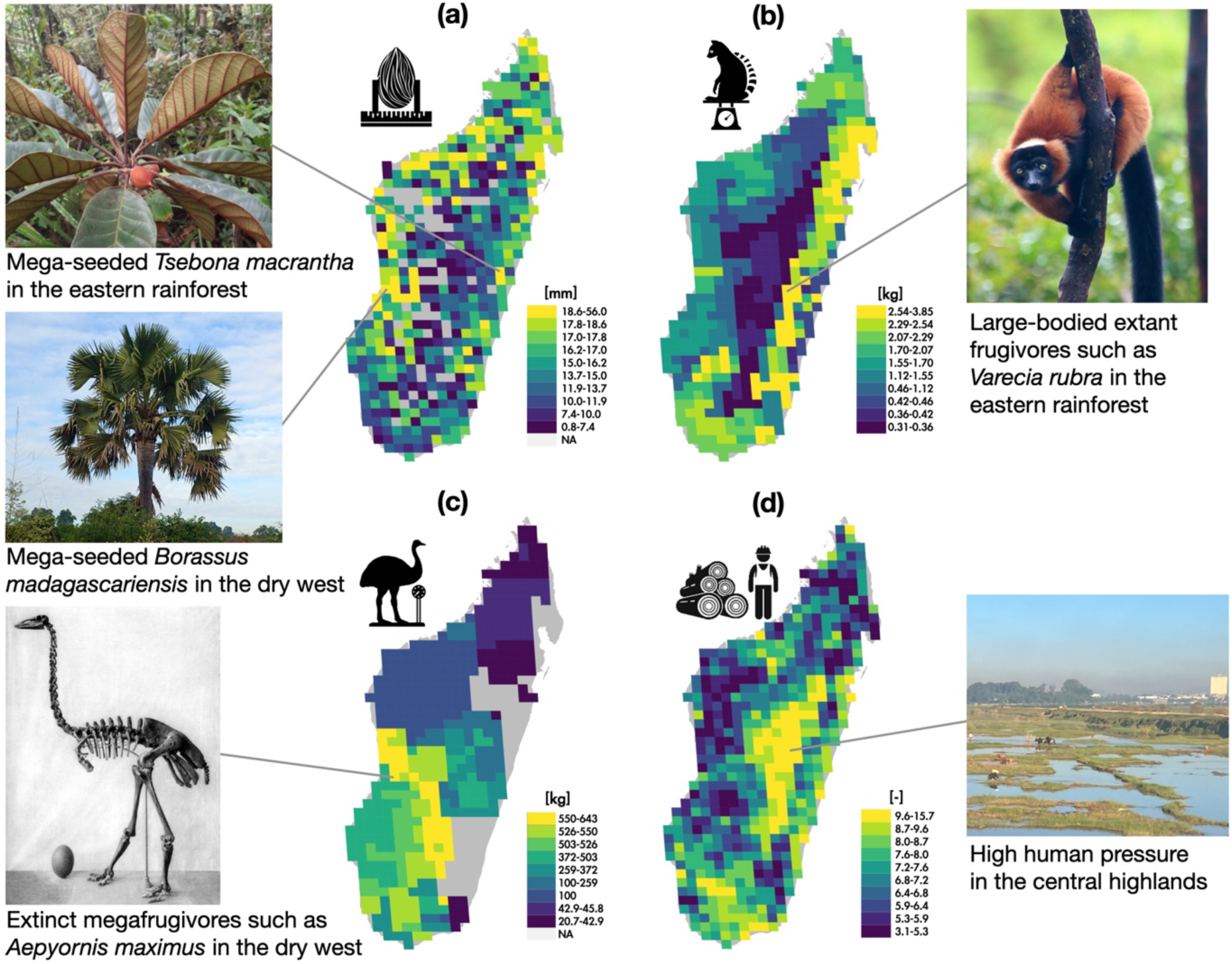
Distribution of seed size, extant and extinct frugivore body mass, and human footprint on Madagascar. These four main variables are used in the structural equation models. (a) 95^th^ percentile maximum plant seed width in each assemblage; (b) 95^th^ percentile maximum body mass of extant frugivores in each assemblage; (c) 95^th^ percentile maximum body mass of extinct frugivores in each assemblage; (d) human footprint of 2009 from Venter et al. (2016). Color gradients of all variables are determined using quantile classification of unprocessed original values into ten classes. Photos of representative species with the largest seeds, largest frugivores, and area with the highest human footprint are shown alongside their locations on the maps. Photo credit: Aina Razanatsima (left top, downloaded from Madagascar Catalogue Image Gallery (http://legacy.tropicos.org/Image/100560806?projectid=17)); Laura Méndez (left middle); Monnier, Quaternary of Madagascar, 1913, (left bottom, downloaded from https://commons.wikimedia.org/wiki/File:Aepyornis_maximus.jpg); Liuye (right top, downloaded from iNaturalist (https://www.inaturalist.org/observations/249596226)); Yuanshu Pu (right bottom).

### PCA of environmental factors

The first two principal components together accounted for 71.25% variance in all environmental factors across assemblages (Fig. 2). PC1 (39.63% variance) reflected a seasonal drought gradient (dry to wet), i.e., positive values were associated with less climatic water deficit, lower evapotranspiration, lower temperature in the warmest months, more precipitation in the driest months, lower precipitation seasonality, and fewer dry months. PC2 (31.62% variance) reflected a gradient from central highland savannas to lowland environments, i.e., positive values were associated with lower temperature seasonality, higher temperature in the coldest months, more annual precipitation, narrower temperature ranges, and lower altitude. By using these two PC axes in the SEMs, we accounted for effects of environmental variation on our focal variables (i.e., plant seed width, extant and extinct frugivore body mass and human footprint).

**Figure 2:**
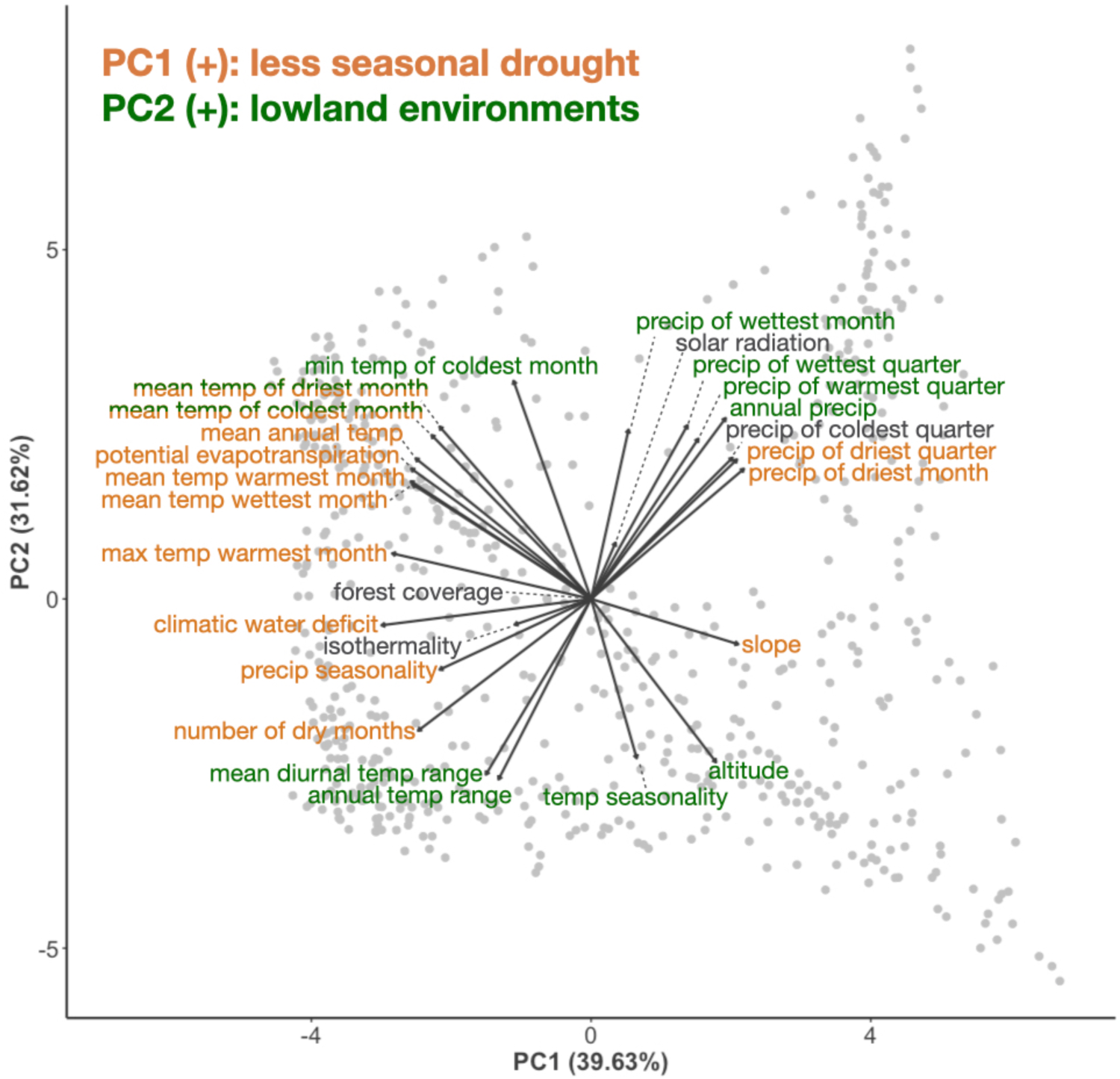
Principal Component Analysis (PCA) of environmental variables across assemblages in Madagascar. Distribution of the first two principal components (PC) are shown. Each dot in the scatter plot corresponds to one of 649 assemblages. Each arrow corresponds to the loadings of an environmental variable. The environmental variables with the highest contributions to PC1 (loadings larger than 0.2) are highlighted in orange, as is the summary of their associated environmental gradient. Similarly, the highest contributing variables (loadings larger than 0.2) and associated environmental gradient for PC2 are highlighted in green.

### Predictors of maximum seed width

In the full SEM model, we found significant effects of all focal predictors on maximum seed width (Fig. 3a, see full SEM fit indices in Table S5). Specifically, human footprint had a direct negative effect on maximum seed width and an indirect negative effect via frugivore downsizing, i.e., decreasing maximum body mass of extant frugivores. In addition, maximum body masses of both extant and extinct frugivores had relatively weak but significant positive effects on maximum seed widths, indicating that seeds are larger in assemblages with larger extant or extinct frugivores. Environmental factors influenced all variables. Specifically, larger seeds and larger extant frugivores were found in lowland environments (PC2). Larger extinct megafrugivores were found in areas that nowadays have more seasonal drought (PC1) and in the central highlands (PC2). Areas with less extreme drought (PC1) and central highlands (PC2) had a higher human footprint. Together, 20.7% variation in maximum seed width was explained by the full SEM.

**Figure 3:**
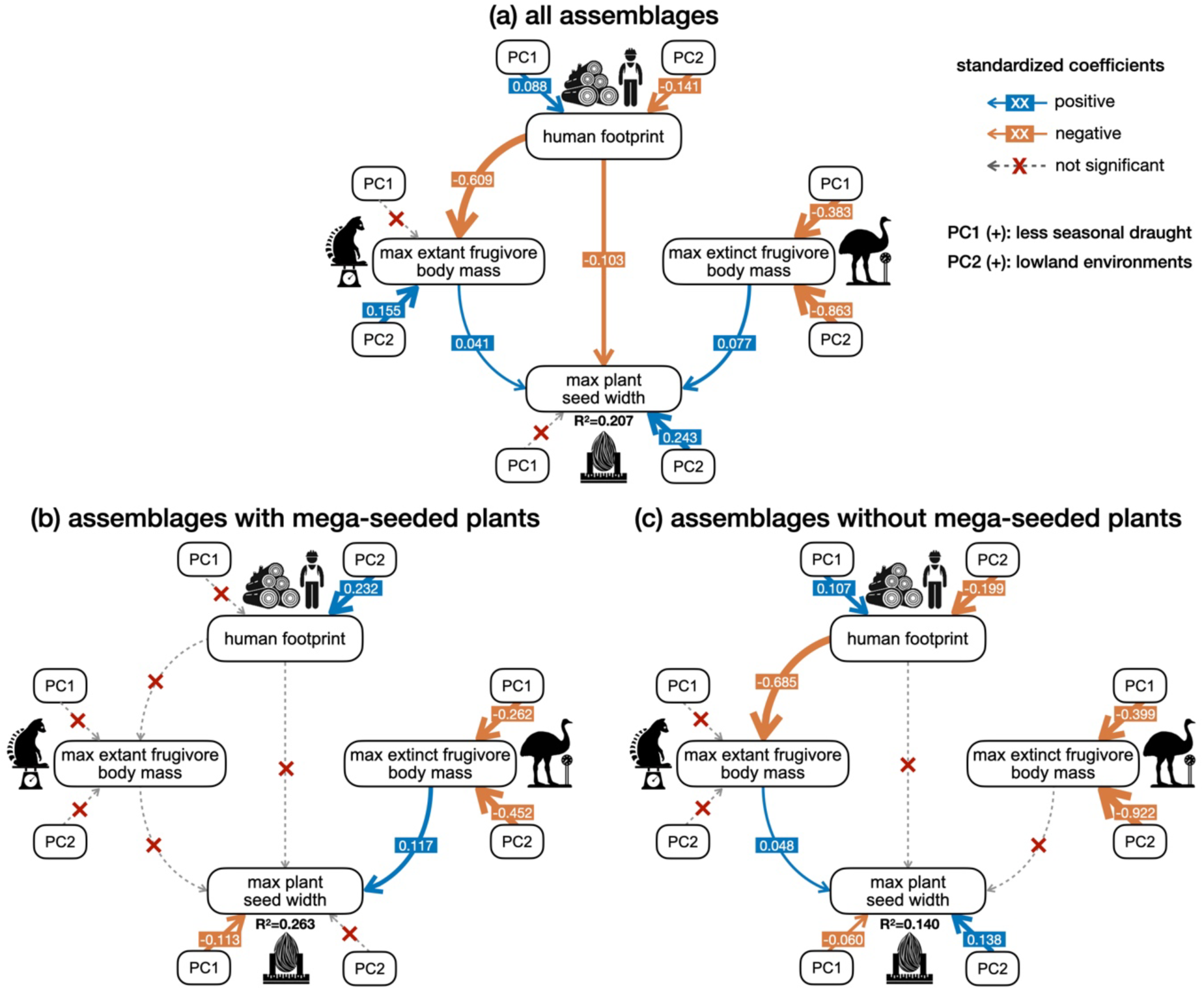
Path diagram showing the relationships between seed width, frugivore body mass, human footprint, and environmental factors in Malagasy plant assemblages. The best fitting structural equation models after model selection are shown, using (a) the Madagascar-wide dataset of 361 assemblages with at least 50% terrestrial area, three endozoochorous plant species, three extant frugivore species and one extinct frugivore species; (b) a subset of 66 assemblages where mega-seeded plants occur (i.e., plants with seeds that are too large to be dispersed by any extant Malagasy frugivore); (c) a subset of 295 assemblages where no mega-seeded plants occur. Effects of the first two principal components of environmental factors on seed width, body masses, and human footprint are shown next to each variable. Covariances allowed in the model are omitted in the figure. The marginal coefficients of determination (R^2^) are given for the focal response variable, max plant seed width. Solid arrows indicate significant effects; dashed arrows with a red cross indicate insignificant effects. The effect sizes of significant standardized coefficients are specified on the arrows, corresponding to the thickness of the arrows (blue = positive effects; orange = negative effects). Max plant seed width = 95^th^ percentile maximum seed width in the assemblage; max extant frugivore body mass = 95^th^ percentile maximum body mass of extant frugivores in the assemblage; max extinct frugivore body mass = 95^th^ percentile maximum body mass of extinct frugivores in the assemblage.

When analysing assemblages with and without mega-seeded plants separately, distinct effects of extant and extinct frugivores were revealed. Specifically, in assemblages with mega-seeded plants (Fig. 3b), maximum seed width was only significantly explained by the maximum body mass of extinct megafrugivores, not extant frugivores. In contrast, in assemblages without mega-seeded plants (Fig. 3c), maximum seed width was only significantly explained by the maximum body mass of extant frugivores, not extinct ones. Additionally, in both subsets, larger seed width was associated with more seasonal drought (PC1). In assemblages without mega-seeds, larger seed width was also associated with lowland environments (PC2). No environmental effects on maximum extant frugivore body masses were detected, but in both subsets, larger maximum body masses of extinct frugivores were found under more seasonal drought (PC1) and in the central highlands (PC2), consistent with the full SEM model. Our predictors explained 26.3% variation in maximum seed width in assemblages with mega-seeded plants, and 14.0% in assemblages without mega-seeded plants.

### Randomisations and spatial autocorrelation

In the 1,000 simulations with randomised maximum seed widths across assemblages, all predictors of seed width became insignificant in ca. 95% of the simulations (see Table S6 for detailed values). Moreover, effect sizes from the empirical data were larger than 95% of the simulated effect sizes for all predictors (Table S6). This emphasizes that relationships revealed from empirical data strongly deviated from a scenario in which maximum seed widths were randomly distributed across assemblages in Madagascar.

Spatial autocorrelation was detected for the residuals in the drivers of maximum seed width using the Madagascar-wide dataset, but not when analysing assemblages with and without mega-seeded plants separately (Table S7). The SAR model for the Madagascar-wide dataset corrected for the spatial autocorrelation in the data, as evidenced by Moran’s I tests (Table S7). Moreover, in the SAR model, all predictors of maximum seed width remained significant (at P < 0.05), but the effect of maximum extant frugivore body mass on maximum seed width was only marginally significant (P = 0.055). All predictors had similar standardized coefficients as in the non-spatial linear regression model (Table S7). This suggests that spatial autocorrelation did not substantially impact our main findings and conclusions.

## Discussion

We found that human footprint had a negative effect on the maximum seed width of an assemblage, both directly, and indirectly by decreasing maximum extant frugivore body mass, supporting the downsizing-effect hypothesis (H1, Fig. 3). Furthermore, our results illustrate that seed widths of anachronistic mega-seeded species are predominantly explained by the past co-occurrence with now extinct megafrugivores, whereas seed widths of smaller-seeded species are shaped by co-occurrence with extant frugivores, thus supporting trait matching and the legacy-effect hypothesis (H2, Fig. 3b, c).

Our results suggest that human footprint could lead to the extirpation of large-seeded plants via two mechanisms. First, via a direct path, human footprint could negatively affect assemblage-level maximum seed width. This indicates that larger-seeded plants may be particularly vulnerable to human impact, or conversely, that smaller-seeded plants are favoured under human footprint. In both cases, it leads to decreases in maximum seed sizes, thus ‘downsizing’ of seeds in assemblages. Indeed, human actions, especially agriculture, overexploitation, and system modifications, are the greatest threats to Malagasy plant biodiversity (Ralimanana et al., 2022). As seed size is correlated to plant size (Leishman et al., 1995; Moles et al., 2005), large trees with large seeds are especially targeted for human utilities such as firewood and construction (Ingram et al., 2005). Moreover, larger-seeded plants are usually characterized by a slower life-history strategy, and are therefore at a disadvantage in the early stages of ecosystem succession after disturbance (Moles & Westoby, 2006). Their slower growth and germination rates make them less competitive compared to faster-establishing pioneer species (Dalling & Hubbell, 2002; Foster & Janson, 1985). Indeed, in human-disturbed and fragmented forests in Madagascar, species turnover typically results in a reduction of tree size and the replacement of forest specialist species by grassland generalists (Hending et al., 2023; Irwin et al., 2010).

Second, we detected an indirect effect of human footprint on seed width, via the trait matching rules in frugivory. Human footprint decreases assemblage-level maximum body masses of extant frugivores, whereas maximum body mass of extant frugivores increases maximum seed widths. Our findings are consistent with evidence on the cascading effects of human-driven defaunation and ‘downsizing’ of animal assemblages on plants, vegetation, and forest structure (Young et al., 2016). For example, frugivore defaunation and downsizing lead to changes in functional outcomes of seed dispersal, such as reduced dispersal distances of seeds and reductions in seed numbers and sizes being dispersed (Donoso et al., 2020; Markl et al., 2012; Pérez-Méndez et al., 2016). Furthermore, many studies have revealed short-term recruitment outcomes of frugivore downsizing, for example, increased seedling aggregation and decreased recruitment of large-seeded plants (Harrison et al., 2013; Kurten, 2013; Razafindratsima et al., 2021). Finally, for longer-term outcomes at the community level, defaunation leads to losses in species richness and diversity (Donoso et al., 2020; Kurten, 2013). Our trait-based study adds to this growing literature, by providing evidence of size-selected changes in specific trait structures of plant assemblages driven by frugivore downsizing and dispersal limitation. Such changes can further alter forest structure and ecosystem function. Seed size is allometrically related to many key plant functional traits, such as plant height, diameter at breast height, and wood density (Hawes et al., 2020; Moles et al., 2005). Hence, downsizing of seeds may affect important ecological functions, such as the potential carbon storage capacity of a species (Chave et al., 2005), and this may ultimately have destructive consequences for the carbon storage of tropical forests as a whole (Bello et al., 2015; Gardner et al., 2019).

With overall larger seeds found in assemblages with larger frugivores, trait matching is evidently still present in extant communities in Madagascar. This size coupling between frugivores and fruits/seeds of endozoochorous plants is consistent with results from plant-frugivore ecological networks (Bender et al., 2018; Burns, 2013; Durand-Bessart et al., 2023; Wheelwright, 1985) and continental-level correlations in trait values between plant and frugivore assemblages (Lim et al., 2020; Lord, 2004; Mack, 1993; McFadden et al., 2022; Wölke et al., 2023). Our results show that this global trend is also recovered at smaller spatial extents and at a finer spatial resolution in Madagascar. However, we acknowledge that our model only explained a small proportion (ca. 20%) of the variation in seed size. This may be partly due to a mismatch in spatial resolution between the occurrence-based distribution of seed size and the polygon-based distribution of frugivore body mass, which could weaken the model’s explanatory power. Additionally, incorporating interaction-based data or other key traits such as the degree of frugivory may better capture the complexity of dispersal interactions and explain more variation in seed size distribution. Although the fossil records used to infer past distributions of megafauna may be spatially biased (but see Méndez et al., 2022; Pedrono et al., 2013), we limited our analysis to assemblages that included extinct frugivores, and the comparative body mass distribution pattern within extinct frugivores should thus be less affected by such biases. It is also important to note that our structural equation model does not elucidate the mechanisms by which downsizing of seeds due to dispersal limitation may happen. Possibly, the loss of large-bodied frugivores leads to secondary extinctions, and hence the extirpation of large-seeded species. Alternatively, large-seeded species are able to adapt, by evolving smaller seeds that can still be dispersed by the remnant frugivore community. Such adaptation can happen rapidly, e.g., within decades, as shown for seeds of the palm *Euterpe edulis* in the Atlantic Forest of Brazil (Galetti et al., 2013).

The abiotic environment – as evidenced by the PCA (Fig. 2) – plays an important role in modulating the size-selective changes in trait structures of plant and frugivore assemblages (Fig. 3). For example, we find that larger seeds generally occur in habitats featuring lowland rainforests or more seasonal drought (Fig. 1a, 3). This aligns with several key theories on the evolution and prevalence of large seeds, namely that large seeds increase survival in shaded forest environment (Leishman & Westoby, 1994) and improve drought/stress tolerance (Westoby et al., 1996; Wölke et al., 2023). Similarly, we find that larger extant frugivores occur in humid lowland environments (Fig. 1b, 3), indicating that the present-day largest frugivorous lemurs depend on rainforest habitats. Although current human footprint remains relatively low in these habitats (Fig. 1d, 3), projected forest loss in Madagascar shows that, even in absence of human impact, these rainforest habitats may decline by 62% solely due to climate change (Morelli et al., 2020).

Even though our model predicts downsizing of seeds due to the potential downsizing of extant frugivores, our empirical data illustrates that 15 mega-seeded plant species have persisted in Madagascar since the extinction of their primary seed dispersers – the Quaternary megafrugivores. These mega-seeded plants seem to have persisted in their historical distribution ranges, with larger seeds found in places with larger past megafrugivores (Fig. 3b). For example, in the dry deciduous forest and the succulent woodlands in west Madagascar, species such as *Borassus madagascariensis* (Arecaceae) and *Adansonia suarezensis* (Malvaceae) bear the largest seeds and fruits in Madagascar (Fig. 1a; seed width up to 67.5 mm wide), possibly once dispersed by the co-existing megafauna such as the elephant birds (Fig. 1a, c; Albert-Daviaud et al., 2020; Janzen & Martin, 1982; Méndez et al., 2022). It remains unclear how these plants have persisted. They may have found alternative dispersal agents, such as introduced zebu cattle, humans, or water (Andriantsaralaza et al., 2024; Guimarães et al., 2008; Méndez et al., 2024; Tonos et al., 2024). Hence, dispersal dysfunctionality is not a major threat to these species. Alternatively, these species have persisted because of long life spans and generation times, but experience shrinking range sizes and decreasing effective population sizes. This has been shown in other places as well, such as on the Canary islands (Pérez-Méndez et al., 2016) and in South America (Doughty et al., 2016; Petrocelli et al., 2024). Possibly, many mega-seeded species have already disappeared since the extinction of the megafauna, although direct evidence from fossil records is scarce, especially in rainforests. At a broader time and spatial scale, increased extinction rates of mega-seeded palms and transition rates to smaller seeds in the Quaternary (last 2.6 million years) hint that downsizing of seeds across assemblages may already be in an progressing stage (Onstein et al., 2018).

In conclusion, we use a causal framework to show direct and indirect downsizing effects of human footprint on seed size in Madagascar. This result emphasizes the urgent need for conservation efforts in areas with increasing human footprint (Semper-Pascual et al., 2023) and that conservation prioritisations should take species interactions and trait-based ecology into consideration (Falcón & Hansen, 2018; Ganzhorn et al., 2024). Furthermore, the distribution of mega-seeded plants holds exclusive signatures of past co-occurring but now extinct megafaunal frugivores, evidencing how current biodiversity patterns cannot be fully understood without considering the (recent) past (Méndez et al., 2022, 2024; Xu et al., 2023). To forecast the future of plant assemblages under continuing frugivore downsizing, future research is needed to better understand how remnant mega-seeded species have persisted in ecosystems without their primary seed dispersers.

## Statement of authorship

YP and REO developed the ideas, hypotheses and methodology for the study. YP and AZ collected and curated the data. YP performed the analyses. REO and AZ provided critical comments and interpretations during analyses. YP and REO wrote the original manuscript. All authors reviewed and edited the manuscript. REO supervised the project.

## Data accessibility statement

All species-level raw data, compiled assemblage-level data, and R scripts used for all analyses in this study will be available in the Dryad Digital Repository upon publication in a peer-reviewed journal.

## Acknowledgements

We acknowledge the support of iDiv, funded by the German Research Foundation (DFG FZT 118, Grant 202548816). We are grateful to our colleagues in the Evolution and Adaptation research group at iDiv for their support and suggestions throughout all stages of this study. In particular, we thank Dr. Laura Méndez for her insights from research in Madagascar, and we thank Andressa Cabral for her valuable input on structural equation modelling methods. We thank Dr. Benjamin Rosenbaum from iDiv for his helpful advice on data analysis.

