## Supporting Document for "Legacy of the lost and pressure of the present: Malagasy plant seeds retain megafauna dispersal signatures but downsize under human footprint"

**Supporting Figures:**

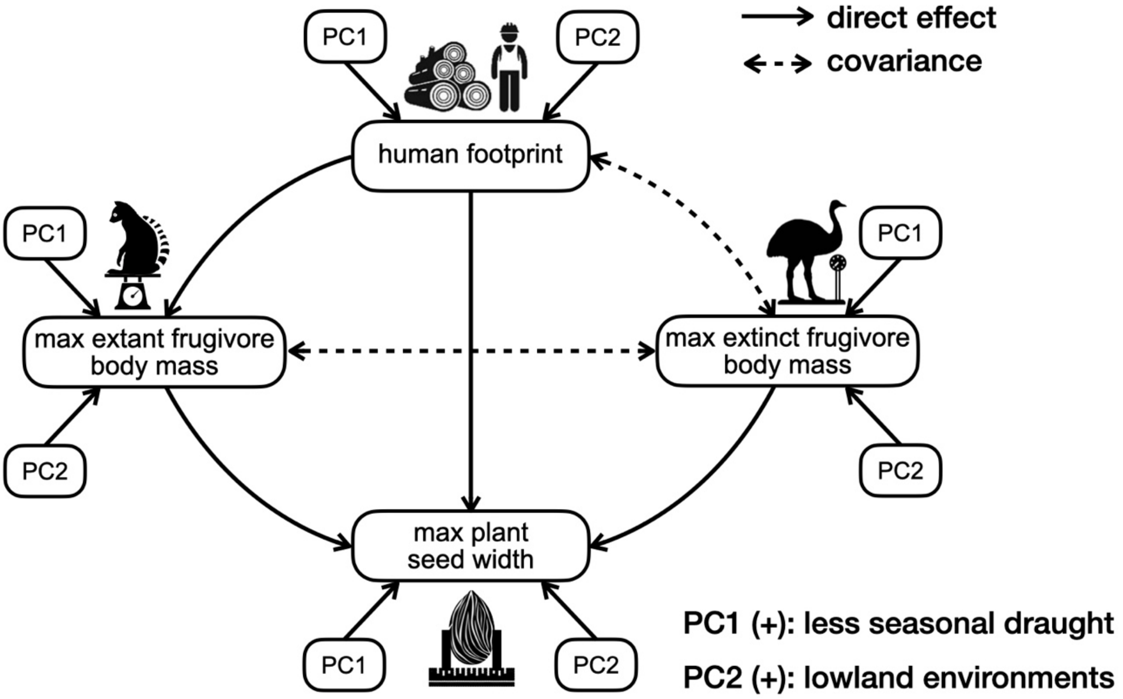


**Figure S1:** ***A priori*** **structural equation model for testing the relationships between seed width, frugivore body mass, human footprint, and environmental factors in Malagasy plant assemblages**. Solid directional arrows indicate direct effects; dashed bidirectional arrows indicate covariance allowed in the model. Effects of the first two principal components of environmental factors on seed width, body mass, and human footprint are shown next to each variable. Max plant seed width = 95$^{th}$ percentile maximum seed width in the assemblage; max extant frugivore body mass = 95$^{th}$ percentile maximum body mass of extant frugivores in the assemblage; max extinct frugivore body mass = 95$^{th}$ percentile maximum body mass of extinct frugivores in the assemblage.

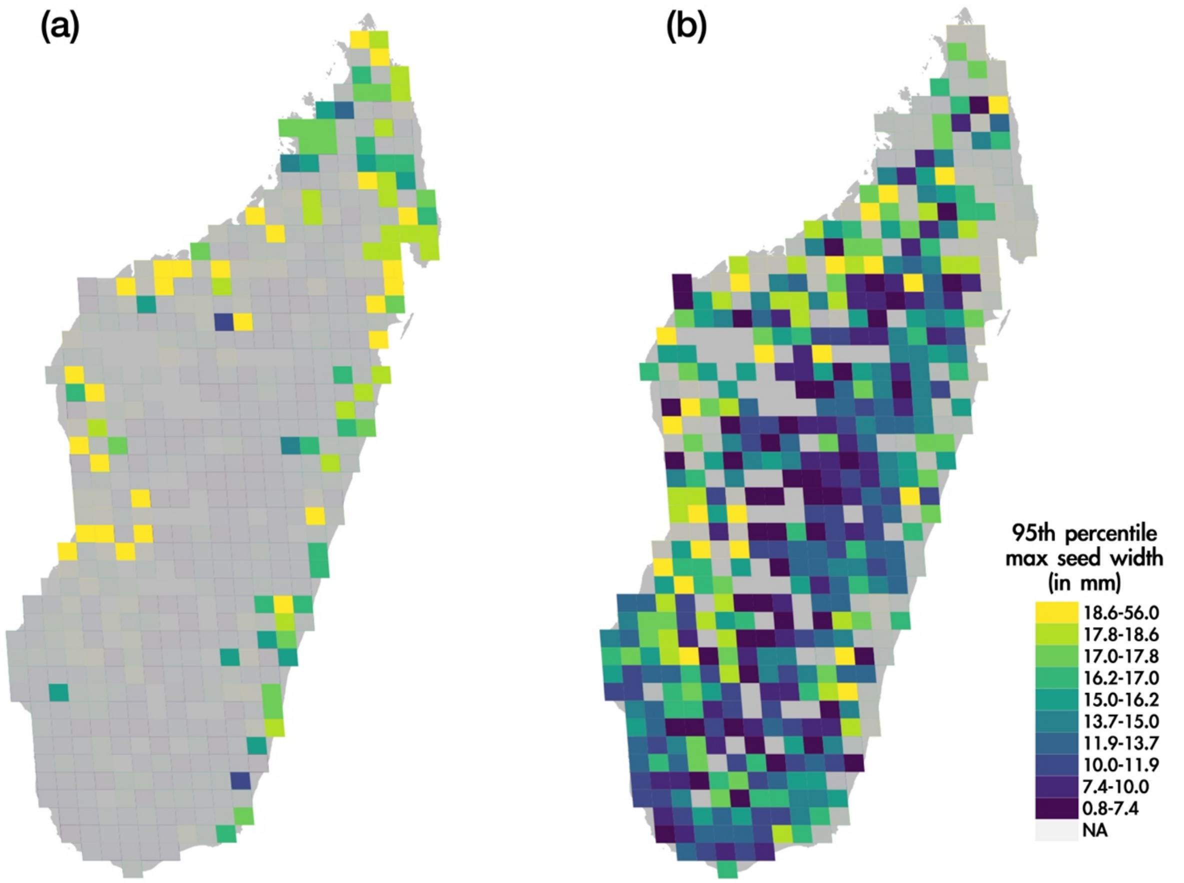


**Figure S2: Distribution maps of the two subsets of Malagasy endozoochorous plant assemblages.** a) 66 assemblages with mega-seeded plant species; b) 295 assemblages without mega-seeded plant species. Species with seeds wider than 24.55mm, the largest seed width dispersed by extant frugivores in Madagascar, are considered as mega-seeded plants. Assemblages that do not belong to each subset are shaded in grey. The colour gradient represents the 95$^{th}$ percentile maximum seed width of each assemblage, and is determined using quantile classification of unprocessed original values into ten classes.

**Supporting tables:**

Table S1-S3 are available in the Dryad Digital Repository; temporary reviewer URL: https://datadryad.org/share/WGJ-Wqr7BlMgKSjBXfN09JwPlopGuMhd8xHClT9NmU4

**Table S1. Taxonomy and fruit/seed traits of 3,241 endozoochorous plant species from Albert-Daviaud et al. (2020).**

See file "Table S1 - 3241 endozoochorous plant species and fruit/seed traits.xlsx".

Column 1-3: species taxonomy

Column 4: whether there's filtered occurrence data from Ralimanana et al. (2022)

Column 5-7: original fruit and seed traits data collected by Albert-Daviaud et al. (2020)

Column 8: seed width including imputed values from Albert-Daviaud et al. (2020)

**Table S2. Taxonomy, frugivory, and body masses of 48 extant frugivores and 15 extinct megafrugivores.**

See file "Table S2 - frugivores species, frugivory and body masses.xlsx".

Sheet 1: 48 extant frugivores

- Column 1-5: species taxonomy
- Column 6: percentage of fruits in the diet of the species
- Column 7: reference for the source of frugivory data
- Column 8: body mass (in kg), averaged across available data from Wilman et al. (2014), Galán-Acedo et al. (2019) and Razafindratsima et al. (2018)

Sheet 2: 15 extinct megafrugivores

- Column 1-2: species taxonomy
- Column 3: body mass (in kg)
- Column 4: reference for the source of body mass data

**Table S3. Species richness, largest seed widths, largest body masses, human footprint, and environmental data in 649 assemblages of plants and frugivores across Madagascar.**

See file "Table S3 - assemblage-level data.xlsx".

Column 1: Assemblage ID

Column 2-3: Coordinates of the assemblage

Column 4: percentage of terrestrial area

Column 5: species richness of endozoochorous plant species

Column 6: 95$^{th}$ percentile maximum plant seed width (in mm, log-transformed)

Column 7: species richness of extant frugivore species

Column 8: 95$^{th}$ percentile maximum body mass of extant frugivores (in kg, log-transformed)

Column 9: species richness of extinct frugivore species

Column 10: 95$^{th}$ percentile maximum body mass of extinct frugivores (in kg, log-transformed)

Column 11: averaged human footprint value (2009), adapted from Venter et al. (2016)

Column 12-14: the first two principal components of the principal component analysis (PCA) of 26 abiotic variables (for a complete overview of variables, see Table S4)

Column 15: whether at least one mega-seeded plant species occurs in the assemblage

**Table S4. List of 26 environmental variables used in this study (source: MadaClim (https://madaclim.cirad.fr/)**

| Category | Variable |
|---|---|
| Temperature | Annual mean temperature (°C) |
| | Mean diurnal range (mean of monthly (max temp - min temp)) (°C) |
| | Temperature annual range (BIO5-BIO6) (°C) |
| | Isothermality (mean diurnal range/temperature annual range x 100) (-) |
| | Temperature seasonality (standard deviation x 100) (°C) |
| | Max temperature of warmest month (°C) |
| | Min temperature of coldest month (°C) |
| | Mean temperature of wettest quarter (°C) |
| | Mean temperature of driest quarter (°C) |
| | Mean temperature of warmest quarter (°C) |
| | Mean temperature of coldest quarter (°C) |
| Precipitation | Annual total precipitation (mm) |
| | Precipitation seasonality (coefficient of variation) (-) |
| | Total precipitation of wettest month (mm) |
| | Total precipitation of driest month (mm) |
| | Total precipitation of wettest quarter (mm) |
| | Total precipitation of driest quarter (mm) |
| | Total precipitation of warmest quarter (mm) |
| | Total precipitation of coldest quarter (mm) |
| Water deficiency | Annual potential evapotranspiration (mm) |
| | Annual climatic water deficit (mm) |
| | Number of dry months in the year (-) |
| Geographic topology | Altitude (m) |
| | Slope (in degree) |
| - | Solar radiation (Wh.m-2.day-1) |
| - | Percentage of forest cover for the year 2010 (%) |

**Table S5. Summary of structural equation modelling final model fit indices, using the full Madagascar-wide dataset (all 361 assemblages), the subset of 66 assemblages with mega-seeded plants, and the subset of 295 assemblages without mega-seeded plants.**

| Indices | Models | | |
|---|---|---|---|
| | All | With megaseeds | Without megaseeds |
| Chi-Square test | 0.100 | 0.319 | 0.243 |
| RMSEA | 0.052 | 0.051 | 0.035 |
| CFI | 0.990 | 0.981 | 0.995 |

Note: The p-value of model Chi-Square test, the root mean square error of approximation (RMSEA) and the comparative fit index (CFI) are shown for the best fitted models using each dataset.

**Table S6. Results of randomization simulations, using the full Madagascar-wide dataset (all 361 assemblages), the subset of 66 assemblages with mega-seeded plants, and the subset of 295 assemblages without mega-seeded plants.**

| Dataset | Effect | Frequency of significant effect in simulation | Observed effect size | 95% confidence interval of effect size |
|---|---|---|---|---|
| whole Madagascar | max plant seed width ~ max extant frugivore body mass | 0.050 | 0.04 | [-0.04,0.04] |
| | max plant seed width ~ max extinct frugivore body mass | 0.043 | 0.08 | [-0.05,0.05] |
| | max plant seed width ~ human footprint | 0.055 | -0.10 | [-0.09,0.10] |
| with megaseeds | max plant seed width ~ max extinct frugivore body mass | 0.052 | 0.12 | [-0.08,0.10] |
| without megaseeds | max plant seed width ~ max extant frugivore body mass | 0.048 | 0.05 | [-0.04,0.04] |

Note: Only predictors relevant to our three main hypotheses and significant in the SEM results were tested and shown. Column “Frequency of significant effect in simulation” indicates the percentage of runs out of 1000 simulations in which the effect was significant at $P < 0.05$.

**Table S7. Results of Spatial autocorrelation test and Spatial autoregressive error model (SAR) for maximum seed width, in comparison with non-spatial linear regression model (LM).**

| Dataset | Moran's I of LM residual | Moran's I of SAR residual | Response | Predictors | Effect size in LM | Effect size in SAR |
|---|---|---|---|---|---|---|
| whole Madagascar | 0.085* | -0.003 | max plant seed width | max extant frugivore body mass | 0.041* | 0.038 . |
| | | | | max extinct frugivore body mass | 0.077** | -0.076** |
| | | | | human footprint | -0.103** | 0.098* |
| | | | | PC2 | 0.243** | 0.243** |
| with megaseeds | -0.21 | - | - | - | - | - |
| without megaseeds | 0.064 | - | - | - | - | - |

Note: dataset “whole Madagascar” refers to the full Madagascar-wide dataset (all 361 assemblages), “with megaseeds” refers to the subset of 66 assemblages with mega-seeded plant, and “without megaseeds” refers to the subset of 295 assemblages without mega-seeded plant species. Only significant predictors from the SEM results were tested and shown. ‘.’ indicates P values smaller than 0.06; ‘*’ indicates P values smaller than 0.05; ‘**’ indicates P values smaller than 0.01.
